# Perturbations of the *AMI1* IAM-amidohydrolase expression trigger plant stress responses in *Arabidopsis thaliana*

**DOI:** 10.1101/2020.08.31.275206

**Authors:** Marta-Marina Pérez-Alonso, Paloma Ortiz-García, José Moya-Cuevas, Thomas Lehmann, Beatriz Sánchez-Parra, Robert G. Björk, Sazzad Karim, Mohammad R. Amirjani, Henrik Aronsson, Mark D. Wilkinson, Stephan Pollmann

## Abstract

The evolutionary success of plants relies to a large extent on their extraordinary ability to adapt to changes in their environment. These adaptations require that plants balance their growth with their stress responses. Plant hormones are crucial mediators orchestrating the underlying adaptive processes. However, whether and how the growth-related hormone auxin and the stress-related hormones jasmonic acid (JA), salicylic acid, and abscisic acid (ABA) are coordinated remains largely elusive. Here, we analyze the physiological role of AMIDASE 1 (AMI1) in plant growth and its possible connection to plant adaptations to abiotic stresses. AMI1 contributes to cellular auxin homeostasis by catalyzing the conversion of indole-acetamide into the major plant auxin indole-3-acetic acid. Functional impairment of AMI1 increases the plants’ stress status rendering mutant plants more susceptible to abiotic stresses. Transcriptomic analysis of *ami1* mutants disclosed the reprogramming of a considerable number of stress-related genes, including JA and ABA biosynthesis genes. The *ami1* mutants exhibit only moderately repressed growth, but an enhanced ABA accumulation, which suggests a role for AMI1 in the crosstalk between auxin and ABA. Altogether, our results suggest that AMI1 is involved in coordinating the trade-off between plant growth and stress responses, balancing auxin with ABA homeostasis.

**HIGHLIGHT:** The IAM amidohydrolase AMI1 catalyzes the conversion of IAM into IAA *in vivo*. Expression of *AMI1* is specifically repressed by osmotic stress conditions, which triggers ABA biosynthesis through the induction of *NCED3*, thereby linking auxin homeostasis with plant stress responses.

## INTRODUCTION

A constantly changing and often adverse environment represents a steady challenge to plants. These challenges include various biotic and abiotic stresses, like pathogen infection, herbivory, high salinity, drought, or temperature changes (Spoel and Dong, 2008; Verma *et al.*, 2016). In order to survive and secure their reproduction, plants often restrict their growth and development under adverse conditions, because energy resources are limited, and their allocation must be tightly balanced to meet the requirements of both growth and adaptive stress responses. To keep track with specific environmental demands, plants have evolved a complex hormone-based network to control their development in response to given changes in their surroundings. Using this system, they can shape their body plan and optimize their metabolism in accordance with prevailing environmental conditions.

Key determinants of this regulatory network are a limited number of signaling molecules, referred to as plant hormones. They orchestrate plant growth and development mainly by taking influence on the transcriptome of responding cells and organs, respectively. Plant hormones act in a combinatorial manner to produce a large number of different responses that are not only dependent on the perceived stimulus, but also on the specific properties of the responding tissue (Depuydt and Hardtke, 2011; Voß *et al.*, 2014). There is mounting evidence for the involvement of auxin in the trade-off between growth and defense. For example, the transcription factor MYC2, which controls the expression of jasmonic acid (JA) responsive genes (Kazan and Manners, 2013), has been reported to negatively regulate the expression of *PLETHORA* (*PLT1* and *PLT2*) transcription factors, which control stem cell development and auxin biosynthesis in roots (Pinon *et al.*, 2013). Other studies also revealed that auxin production is increased by JA through the induction of *ANTHANILATE SYNTHASE 1* (*ASA1*) and a small number of *YUCCA* genes in certain plant tissues (Hentrich *et al.*, 2013; Sun *et al.*, 2009).

Plants also balance their growth with responses to abiotic stresses (Verma *et al.*, 2016; Zhu, 2002). When exposed to drought or high salinity, two major abiotic stress factors in the field, plants reduce their growth rate, while stress-resistance mechanisms are initiated. Through the flexible reprogramming of their gene regulatory networks, plants strive for attaining the best suited phenotype for the prevailing stress condition (Claeys and Inzé, 2013; Julkowska and Testerink, 2015). Brassinosteroid signaling pathways are reported to contribute to the control of plant growth under drought or starvation stress by regulating autophagy (Nolan *et al.*, 2017). However, other plant hormones including auxin and abscisic acid (ABA) are also assumed to contribute to the coordination of the growth–stress response trade-off. Auxin is known to be the crucial determinant for plant growth (Davies, 2010), whereas ABA is reported to operate as a stress hormone in responses to abiotic stimuli (Gomez-Cadenas *et al.*, 2015; Ma *et al.*, 2018).

Under optimal conditions, plants usually grow rapidly with their energy resources mainly dedicated to primary growth, including root system- and leaf expansion, shoot elongation, and reproduction. In this scenario, auxin is pivotal because it governs virtually all aspects of plant growth and development through the promotion of cell elongation, expansion, and differentiation (Vanneste and Friml, 2009). To ensure optimal plant growth, auxin homeostasis needs to be tightly regulated (Ljung, 2013). In order to control cellular auxin contents, plants possess a series of different biochemical and biological tools, including *de novo*-biosynthesis, inactivation through conjugation, sequestration, and degradation (Brumos *et al.*, 2018; Ruiz Rosquete *et al.*, 2012). In contrast, when plants are subjected to biotic and abiotic stresses, they respond with the adjustment of their growth program, which in many cases results in reduced growth rates and premature phase transition from vegetative to reproductive growth (Perdomo *et al.*, 2015; Silva *et al.*, 2013; Suzuki *et al.*, 2014). JA and ABA, two well-characterized plant stress hormones, play crucial roles in plant responses to biotic and abiotic stress factors. However, an underlying mechanism by which those two phytohormones are connected with cellular auxin levels remains largely elusive.

Mainly based on *in vitro-*evidences, AMI1 has been suggested to act in a side pathway of indole-3-acetic acid (IAA) biosynthesis (**Fig. 1**), converting indole-3-acetamide (IAM) into IAA (Nemoto *et al.*, 2009; Neu *et al.*, 2007; Pollmann *et al.*, 2003; Sánchez-Parra *et al.*, 2014). In Arabidopsis, the majority of IAM (95%) has been reported to derive from the precursor indole-3-acetaldoxime (IAOx) (Sugawara *et al.*, 2009). The enzyme responsible for the biochemical conversion, however, has yet to be identified. IAOx depleted mutant plants, i.e. *cyp79b2 cyp79b3*, neither show significant alterations of free auxin contents under standard conditions nor transcriptional induction of other auxin biosynthesis pathway components to compensate the loss of IAOx-dependent pathway(s) (Lehmann *et al.*, 2017; Zhao *et al.*, 2002). Together, these findings argue against a contribution of AMI1 in general auxin biosynthesis. Here, we report the functional characterization of AMIDASE 1 (AMI1) *in vivo*. The comprehensive analysis of *AMI1* gain- and loss-of-function mutants provided indication for an involvement of AMI1 in auxin homeostasis. A loss of *AMI1* results in increased IAM levels and moderately reduced IAA contents, whereas conditional overexpression of *AMI1* (AMI1ind) had the opposite effect. The examined *ami1* mutant alleles, *ami1-1* and *ami1-2*, show moderate growth reductions, while independent AMI1ind lines are characterized by auxin overproduction-related phenotypes. Comprehensive microarray analyses comparing *AMI1* mutants with wild-type plants revealed a tight connection of *AMI1* with various biotic and abiotic stresses, as numerous stress marker genes appeared to be differentially expressed. Abiotic stress treatments confirmed that *ami1* mutants are more susceptible towards osmotic stress. In summary, our results suggest that AMI1 is involved in the coordination of the trade-off between growth and stress resistance, rather than in the *de novo*-biosynthesis of auxin.

**Figure 1.**
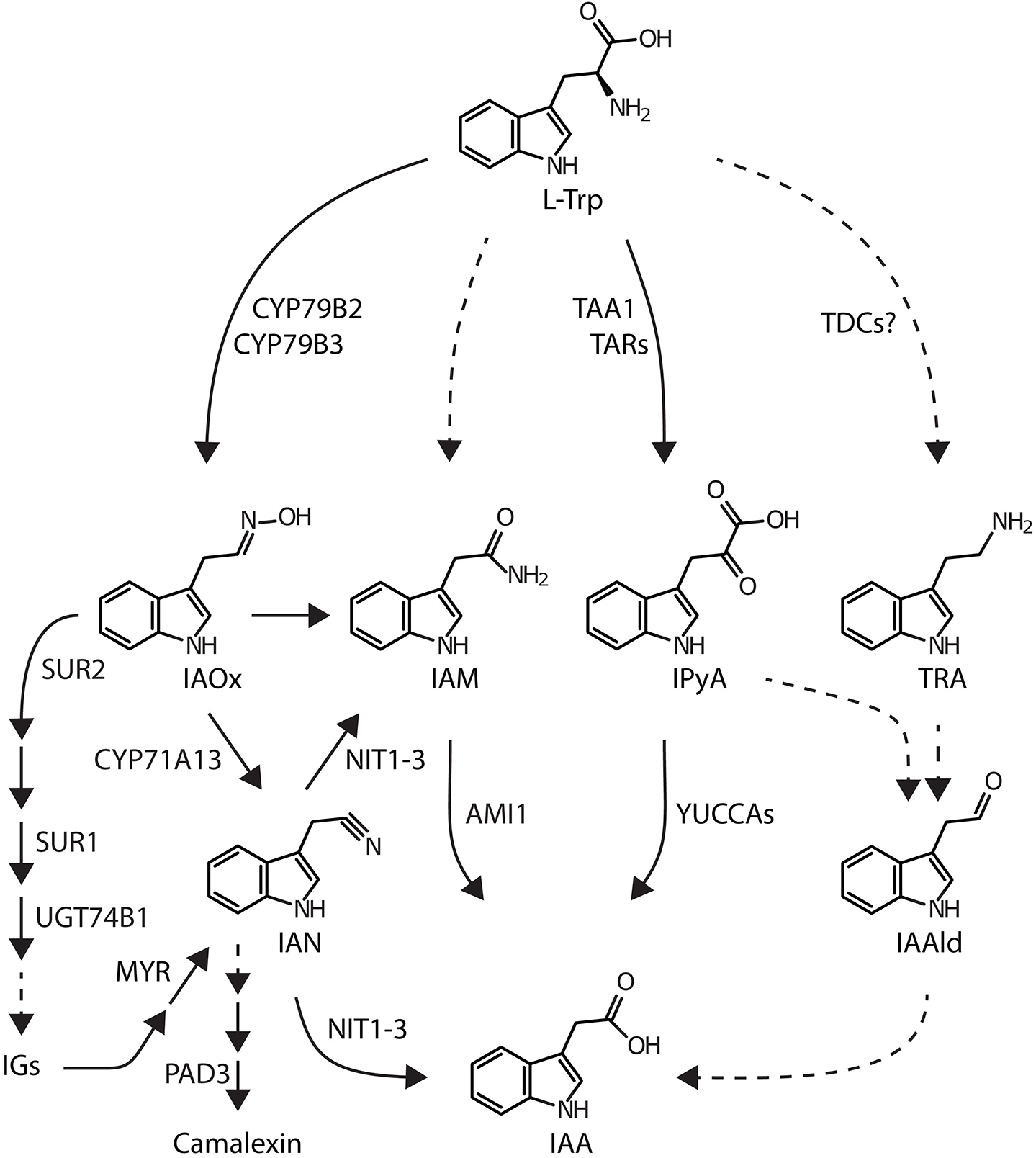
Proposed anabolic routes for IAA biosynthesis in *Arabidopsis thaliana*. Dashed lines represent assumed reaction steps for which the corresponding genes/enzymes have not yet been identified. Metabolites and enzymes are abbreviated as follows: AMI1, AMIDASE 1; CYP71A13, CYTOCHROME P450 MONOOXYGENASE 71A13; CYP79B2/B3, CYTOCHROME P450 MONOOXYGENASE 79B2/B3; IAA, indole-3-acetic acid; IAAld, indole-3-acetaldehyde; IAM, indole-3-acetamide; IAN, indole-3-acetonitrile; IAOx, indole-3-acetaldoxime; IGs, indole glucosinolates; IPyA, indole-3-pyruvic acid; L-Trp, L-tryptophan; MYR, MYROSINASE; NIT1-3, NITRILASE 1-3; PAD3 PHYTOALEXIN DEFICIENT 3 (CYTOCHROME P450 MONOOXYGENASE 71B15); SUR1, SUPERROOT 1 (S-ALKYL-THIOHYDROXYMATE LYASE); SUR2, SUPERROOT 2 (CYTOCHROME P450 MONOOXYGENASE 83B1); TAA1, TRYPTOPHAN AMINOTRANSFERASE OF ARABIDOPSIS 1; TAR, TRYPTOPHAN AMINOTRANSFERASE RELATED; TDC, TRYPTOPHAN DECARBOXYLASE; TRA, tryptamine; UGT74B1, UDP-GLUCOSYL TRANSFERASE 74B1.

## MATERIALS AND METHODS

### Plant material and plant growth conditions

The experiments were carried out using *Arabidopsis thaliana* (L.) Heynh. ecotype Col-0 (from Nottingham Arabidopsis Stock Centre (NASC), stock N1092). Seeds for *35S::tms2* (N6265), and the T-DNA insertion lines for *AMI1* (At1g08980; *ami1-2* (SALK_019823C)) were obtained from the NASC, or kindly provided by Dr. Henrik Aronsson, *ami1-1* (Aronsson *et al.*, 2007). T-DNA mutants were genotyped according to (Alonso *et al.*, 2003). Primers for genotyping are given in **Table S1**. If not stated otherwise, seedlings were raised under sterile conditions on solidified ½ MS media containing 1% (w/v) sucrose in Petri dishes (Murashige and Skoog, 1962). Plantlets were kept under constant environmental short day (SD) conditions (8 h light at 24 °C, 16 h darkness at 20 °C, photosynthetically active radiation 105 μmol photons m^−2^ s^−1^) for 2 to 3 weeks. If older plant material was used, the plant organs were harvested from plants grown in a greenhouse on a mixture of soil and sand (2:1) for 4-6 weeks in SD. Thereafter, plants were transferred to long day (LD) conditions (16 h photoperiod). The greenhouse was maintained under constant climatic conditions, 22 to 24 °C during daytime and 18 to 20 °C overnight. The photosynthetically active radiation was no less than 150 μmol photons m^−2^ s^−1^. Expression of the transgene in AMI1ind lines was induced by either 10 μM β-estradiol added to the growth media or by the administration of 50 μM β-estradiol through irrigation.

### Generation of transgenic plants

Transgenic plants conditionally overexpressing AMI1 with a N-terminal *c-myc* tag (AMI1ind) and containing AMI1 promoter reporter constructs (*pAMI1::GUS*), respectively, were generated as described in **Protocol S1**.

### Transcriptomics

Total RNA was extracted from 100 mg plant tissue as previously described (Oñate-Sánchez and Vicente-Carbajosa, 2008). Labeling and hybridization of cDNA libraries from *ami1-2*, AMI1ind-2, and Col-0 to ATH1 DNA chips (Affymetrix) was performed by the CNB Genomics Service (Madrid, Spain). For each genotype three biological replicates were processed. Variation between replicates was accounted for by using the LIMMA model (Ritchie *et al.*, 2015). Differentially expressed genes were identified by the modified *t*-test implemented in the LIMMA package. A Benjamini-Hochberg correction was used to adjust for multiple testing. An adjusted *p*-value < 0.05 and fold-change ≥ 1.5 were arbitrarily chosen to select 378 and 878 differentially expressed genes (DEGs) in *ami1-2* and AMI1ind-2, respectively, relative to wild type (**Table S2**). The Gene Set Enrichment Analysis (GSEA) used a hypergeometric test with a Benjamini-Hochberg FDR correction and a post-correction selection significance level of *p* < 0.05 (Maere *et al.*, 2005). Complementing the GSEA, Parametric Analysis of Gene Set Enrichment (PAGE) analyses were executed to determine GO terms enriched in the de-regulated gene groups. The PAGE method is statistically more sensitive and accounts for both the number of genes and their respective expression patterns, employing a Benjamini-Hochberg (FDR) multi-test adjustment and a significance of *p* < 0.05 (Kim and Volsky, 2005).

Selected transcripts were validated in independent experiments by qRT-PCR. For each genotype and condition, total RNA from three independent biological replicates was harvested and analyzed in triplicate (technical replicates). First-strand synthesis was performed according to the supplier’s instructions, using M-MLV reverse transcriptase and oligo(dT)_15_ primer (Promega). Two ng of cDNA was used as template in qPCRs, which were performed according to the manufacturer’s instructions using the FastStart SYBR Green Master solution (Roche Diagnostics) on a Lightcycler 480 Real-Time PCR system (Roche Diagnostics). Relative quantification of expression was calculated after data analysis by the Lightcycler 480 software (Roche Diagnostics), using the comparative 2^−ΔΔC^T method (Livak and Schmittgen, 2001) with *APT1* (At1g27450) and *UBQ10* (At4g05320) as reference genes (Czechowski *et al.*, 2005; Jost *et al.*, 2007). See **Table S1** for primer sequences.

### Assay for ß-glucuronidase activity

Histochemical ß-glucuronidase (GUS) assays were performed as described by Jefferson *et al.* (1987). Seedlings and detached plant parts, respectively, were infiltrated with a GUS staining solution, consisting of 1 mM 5-bromo-4-chloro-3-indoxyl-ß-D-glucuronic acid (X-Gluc), 50 mM potassium phosphate buffer, pH 7.0, 1 mM EDTA, 0.1% Triton X-100, and incubated for 16 h at RT. Chlorophyll was removed by washing with an ethanol series (30%, 70%, and 96% (v/v)), and the tissue was rehydrated in water for photography.

### Amidase activity assay

The IAM hydrolase activity in plants was determined as previously described (Neu *et al.*, 2007). In brief, crude extracts from 2-week-old seedlings were prepared. One hundred milligram of plant tissue was shock-frozen in liquid nitrogen. The tissue was disrupted using micro-pestles. As the material started to thaw, buffer (50 mM HEPES, pH 8.5, 200 mM sucrose, 3 mM EDTA, 3 mM DTT, 5% (w/v) insoluble polyvinylpyrrolidone) was added in a ratio of 1:3 (w/v). The suspension was homogenized until no more tissue fragments were visible. Cell debris and insoluble matters were collected by centrifugation (16.000 *g*, 15 min, 4 °C) and the supernatant was transferred to fresh tubes. Protein concentrations were determined according to Bradford (1976), using serum albumin as a protein standard. Aliquots containing 200 μg of protein were incubated with 5 mM IAM in a total volume of 300 μl of 50 mM potassium phosphate buffer (pH 7.5) for 3 h at 30 °C. Finally, the amount of produced IAA was quantified by RP-HPLC. The data were analyzed using Student’s *t*-test to compare two means. Statistical analyses were conducted using PRISM version 7.0a (GraphPad Software).

### IAA and IAM quantification

The LC-MS analysis of endogenous IAA and IAM was carried out according to Novák *et al.* (2012). For each sample 100 mg of plant material were harvested and immediately frozen in liquid nitrogen. Each independent experiment used, at least, three biological replicates. Sample handling times were kept as short as possible to prevent distortion of the IAA pool through the autocatalytic decay of precursor molecules (Gélinas-Marion *et al.*, 2020). More detailed information on the applied LC-MS settings can be found in **Protocol S1**.

### ABA quantification

The ABA contents in 50 mg of 10 day-old sterilely grown Col-0 and *ami1* seedlings were determined using GC-MS/MS (Bruker Daltonics, Scion-TQ) analysis as previously described (Ramos *et al.*, 2018). Quantification was carried out using stable isotope-labelled [^2^H_6_]-ABA as internal standard. Endogenous hormone contents were calculated from the signal ratio of the unlabeled over the stable isotope-containing mass fragment observed in measurements that have been performed in quintuplicate. The data were analyzed with one-way ANOVA followed by Tukey’s B post-hoc test to allow for comparison among all means. Statistical analyses were conducted using PRISM version 7.0a (GraphPad Software).

### Western blotting

Both SDS-PAGE and amidase immunoblots were carried out as described by Neu *et al.* (2007). A monoclonal α-myc antibody, clone 9.E10 (Evan *et al.*, 1985) was used as first antibody at a final dilution of 65 mg l^−1^. As second antibody, a rabbit-anti-mouse-IgG (Promega) coupled with an alkaline phosphatase was used.

### Biomass and root morphology analysis

For the analysis of the root morphology, five replicate Petri dishes with ½ MS agar medium (0.5% sucrose, w/v), containing 21-26 seedlings, were used for each line. The root system morphology of the seedlings was investigated using the WinRhizo™ image analysis software (Regent Instruments Inc.). The roots were scanned using a 600 dpi resolution STD1600+ scanner (Regent instruments Inc.) while still attached to the dish. The images were analyzed to identify various root system morphology parameters (Björk *et al.*, 2007), namely root length (cm), surface area (cm^2^), number of branches (forks), and number of root tips. The shoot and the root systems were then separated and their dry weight determined using a Sartorius ultra-micro scale (Sartorius GmbH, Göttingen, Germany) after drying the individual parts at 70 °C for at least 48 h. Root-to-shoot ratio (R/S ratio), specific root length (SRL; m g_DW_^−1^ root), specific root area (SRA; m^2^ kg_DW_^−1^ root), specific root tip density (tips g_DW_^−1^ root), root tip density (tips cm^−1^ root), specific branching density (forks g_DW_^−1^ root), and branching density (forks cm^−1^ root) were calculated for each genotype (Björk *et al.*, 2007). Differences in the root architecture were investigated using a MANOVA, which was followed by Tukey’s HSD post hoc test for the genotype. Statistical analyses were conducted using SPSS 18.0 (SPSS Inc.).

## RESULTS

### The *AMI1* gene is generally expressed in proliferating tissues, but repressed during early stages of germination

In our previous work, we performed microscopic experiments using AMI1-GFP fusion proteins and semi-quantitative RT-PCR to investigate the subcellular localization and tissue specific expression of *AMI1*. The experiments revealed a mainly cytoplasmic localization and the expression of *AMI1* in proliferating tissues, such as young leaves and flowers (Pollmann *et al.*, 2006b). Additional qPCR experiments confirmed the expression pattern for *AMI1* on a tissue-specific basis. *AMI1* expression was detected in all analyzed samples. The expression in roots was, however, low compared to the levels in all other tissues. In adult plants, *AMI1* expression was highest in juvenile leaves and petioles; lower transcript levels were detected in stems and in flower buds (**Fig. S1A**).

To enable further examination of *AMI1* expression during plant development, we generated *pAMI1::GUS* promoter reporter lines. Since qPCR data pointed towards high *AMI1* expression in young seedlings, we monitored ß-glucuronidase (GUS) activity in *pAMI1::GUS* lines between 26 h and 72 h after imbibition (**Fig. 2A**). The GUS staining showed an increasing *AMI1* promoter activity with the ongoing of seedling development. Starting after about 48 h in the cotyledons, GUS activity peaked 64 h after germination. The observed promoter activity matches considerably well with the previously determined IAM profile during seed germination and early seedling development (Hoffmann *et al.*, 2010; Pollmann *et al.*, 2002), pointing towards a rapid decline of IAM levels during early seedling development. At later stages, especially in the primary root, the expression slowly decreased, while in root tips a clear GUS activity remained detectable. The qPCR experiment provided further evidence for a substantial *AMI1* expression in leaves, petioles, and flower buds. For this reason, we analyzed *AMI1* promoter activity during later stages of development. In cotyledons, expression was mainly detected in the vascular tissue and on the very tip of the leaf (**Fig. 2B**). In primary leaves, GUS staining was visible in the vascular tissue, at the tip of the leaf, and also in trichomes (**Fig. 2C**). During flower development, at developmental stage 14 (Smyth *et al.*, 1990), strong activity was detected in the petioles, pedicels and receptacles, in the vasculature of the sepals, and at the end of the pistil including the stigma (**Fig. 2D**). Only low activity was visible in the filaments and anthers, whereas the petals showed no staining (**Fig. 2D,E**). In developing siliques, at stage 17, strong promoter activity was observed at the end of the pedicels and on both ends of the silique, whereas nearly no activity was found in the developing seeds and funiculi (**Fig. 2F**). Taken together, these results confirm the expression of *AMI1* in proliferating tissues, including young seedlings and developing flowers.

**Figure 2.**
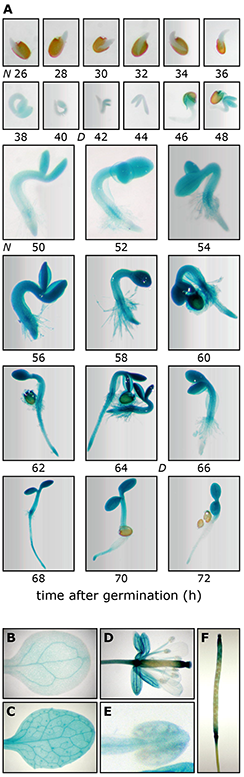
Expression pattern of *AMI1* during plant development. **(A)** GUS staining of *pAMI1::GUS* seedlings 26 to 72 h after imbibition. Plants were grown on ½ MS media under short day conditions (8 h light/16 h darkness) and then stained for reporter activity. The light-dark cycle is indicated by N (night) and D (day), respectively. **(B)** GUS staining of a 5-day-old cotyledon from a *pAMI1::GUS* seedling. Staining was observed at the cotyledon tip and in the vascular tissue. **(C)** GUS staining of a 6-week-old *pAMI1::GUS* leaf. Staining was located in the vascular tissue and at the tip of the leaf. **(D)** GUS staining of a mature flower from a *pAMI1::GUS* plant at developmental stage 12. Note the strong staining in the sepal vasculature, the pedicel, and on both ends of the carpel, including the stigma. **(E)** Magnification of a GUS stained anther from the flower shown in **(D)**. **(F)** GUS staining of a developing silique at stage 17. Strong GUS activity can be observed at the ends of the siliques and at the end of the pedicel. The developmental stages of the flowers were classified according to Smyth et al. (1990).

### High levels of IAM induce *AMI1* gene expression

Numerous previous studies already analyzed the transcriptomic response towards IAA in Arabidopsis. The inspection of publicly available datasets, e.g. GSE631 and GSE42007 (Lewis *et al.*, 2013; Okushima *et al.*, 2005), demonstrated that most auxin homeostasis-related genes show no significant response to IAA (**Table S2**). Consistent over both datasets, only a small number of *Gretchen Hagen 3* (*GH3*) genes appear to be significantly induced by IAA. *GH3* genes encode IAA-amidosynthases that catalyze the conjugation of free IAA to amino acids, thereby physiologically inactivating the plant hormone (Staswick *et al.*, 2005). To investigate the response of *AMI1* towards the exogenous application of IAA and IAM in closer detail, we performed qPCR experiments on 2-week-old seedlings, which were treated for different periods of time with the indoles. Generally confirming the transcriptomic data, the application of IAA (20 μM) triggered only a weak repression (*log2* = −1.3 ± 0.1) of *AMI1* transcription. In contrast, the treatment with 20 μM IAM significantly induced the expression of *AMI1*. The transcriptional level of *AMI1* temporarily increased to reach a *log2* value of 2.9 ± 0.3 at 2h, only to return to a lower level after 3h of IAM treatment (**Fig. 3**). The induction of *AMI1* gene expression by its putative substrate, IAM, suggests a function of AMI1 in the control of cellular IAM levels.

**Figure 3.**
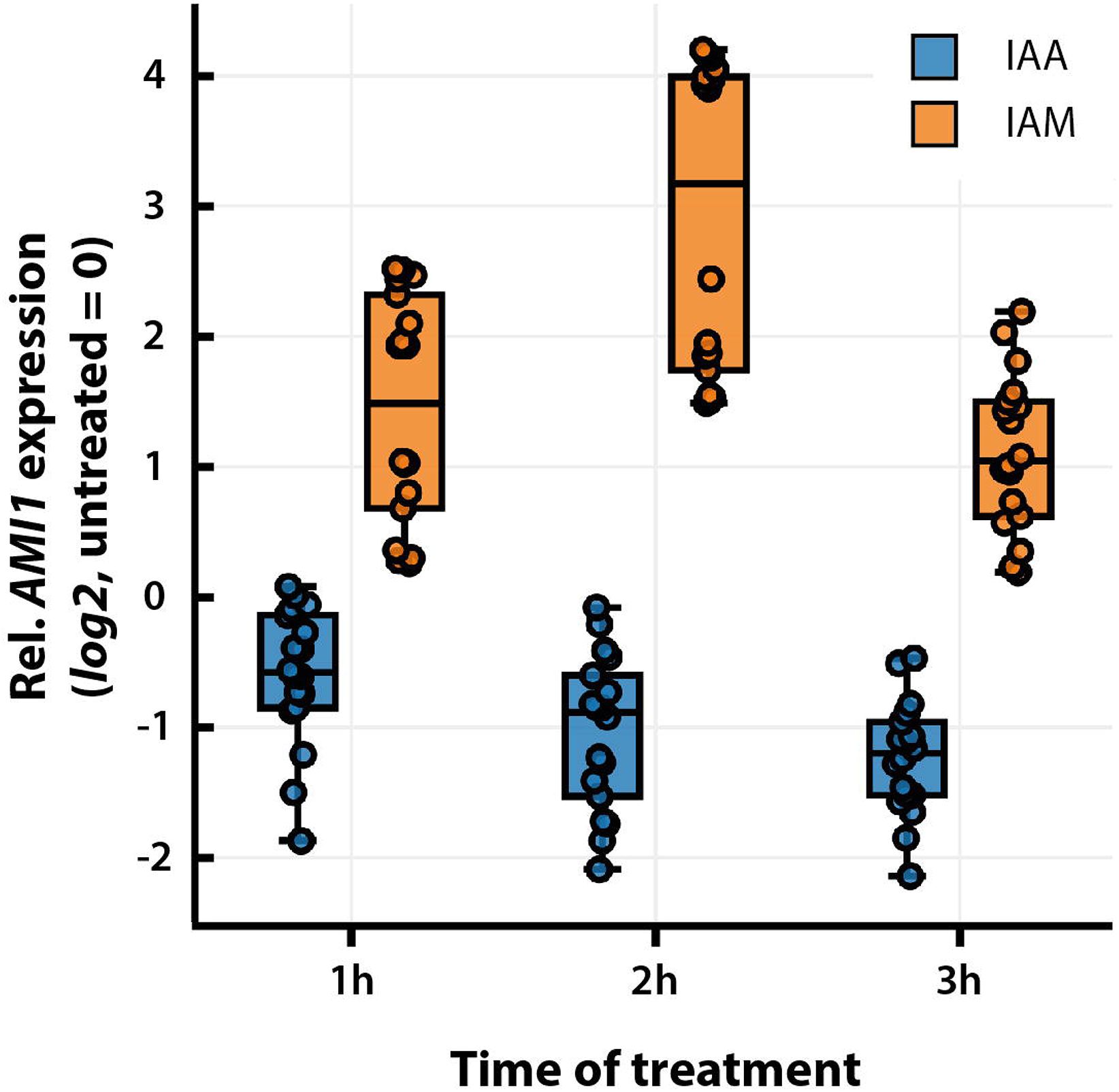
Transcriptional response of *AMI1* towards exogenously applied IAA and IAM. Transcript levels of *AMI1* in treated seedlings were measured by qRT-PCR. One hundred milligram plant tissue was harvested for each sample after 1, 2, and 3 h of treatment and used for RNA extraction. Transcript abundance values are normalized to the geometric means of *APT1*, and *UBQ10* transcripts and given relative to *AMI1* expression levels in mock treated samples. Means are given with their SE (n = 18).

### Perturbations of *AMI1* gene expression alter auxin contents and trigger auxin-related phenotypes

To explore the role of AMI1 in auxin homeostasis, we took a complementary reverse genetics approach. First, we studied two independent mutant alleles for *AMI1*, carrying T-DNA insertions either in the fourth intron (*ami1-1*) (Aronsson *et al.*, 2007) or in the sixth exon (*ami1-2*). The T-DNA insertion sites of *ami1-1* and *ami1-2* were confirmed by sequencing. In case of *ami1-2*, the sequencing of the genomic DNA revealed a tandem T-DNA insertion (**Fig. 4A**). Homozygous plants were isolated and compared to wild-type Arabidopsis (**Fig. 4B**). Both null alleles were devoid of detectable amounts of full-length *AMI1* mRNA (**Fig. S1B**), which could also be validated by qPCR, demonstrating a lack of expression of the second half of the *AMI1* gene, containing the T-DNA insertions (**Fig. S1C**). The results provided evidence that the T-DNA insertions in *ami1-1* and *ami1-2* result in the formation of truncated and likely functionally impaired transcripts.

**Figure 4.**
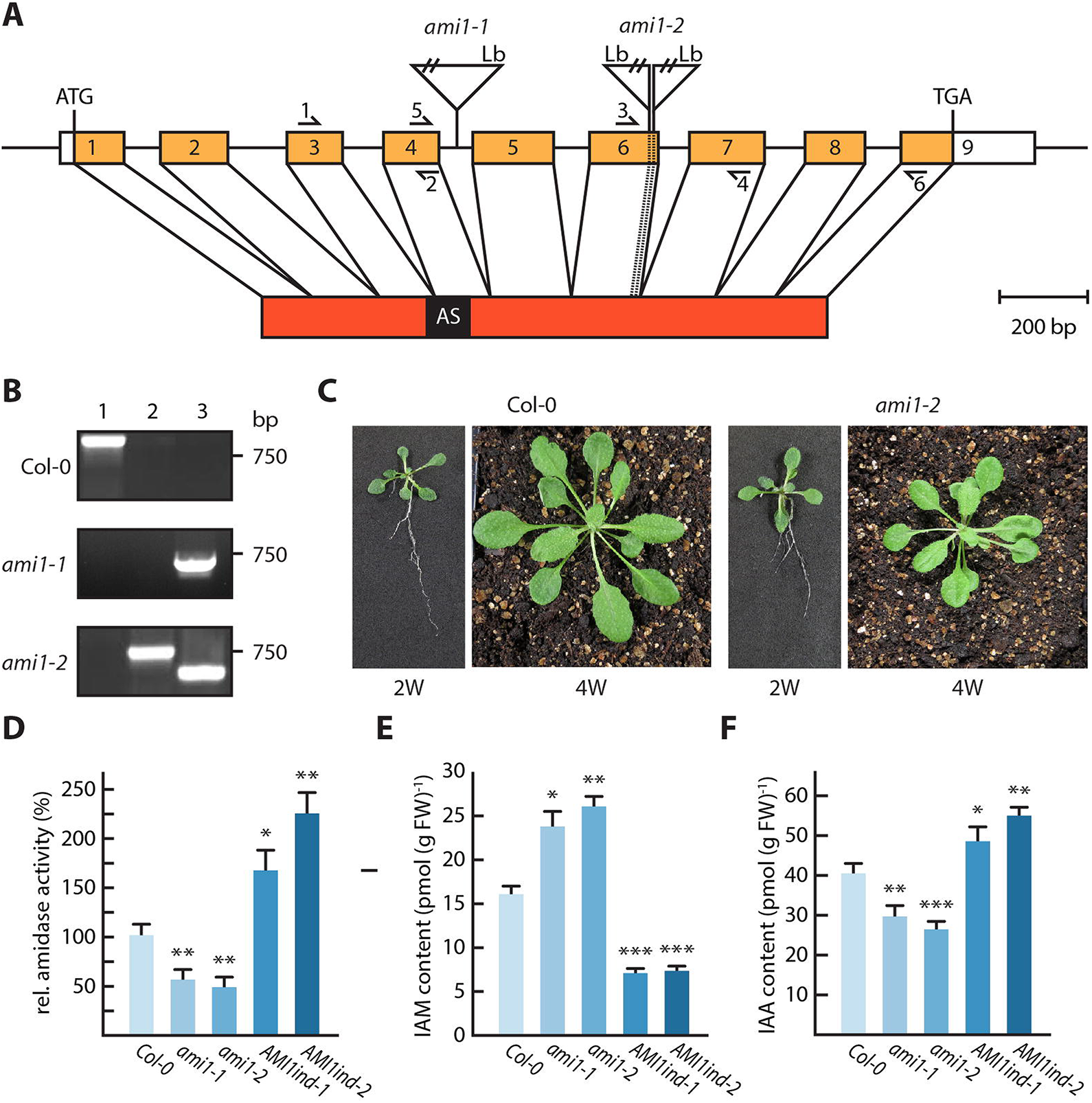
Characterization of *ami1* T-DNA insertion lines. **(A)** Genomic *AMI1* region (At1g08980) showing the exon/intron structure and the T-DNA insertion sites in intron 4 and exon 6. The primer pairs used for either RT-PCR (1/2, 3/4) or qRT-PCR (5/6) are shown as arrows. The resulting cDNA with the fused exons, but without the 5’ and 3’ untranslated regions, is depicted below. Also given is the amidase signature (AS) domain of the derived protein. Insertion sites of the T-DNA in the alleles *ami1-1* and *ami1-2* were analyzed by sequencing. Genomic sequence and cDNA, but not the T-DNA, are drawn to scale. Lb, left boarder; ATG, start codon; TGA, stop codon. **(B)** PCR zygosity analysis of the studied genotypes. Total genomic DNA was extracted from wild type and homozygous T-DNA insertion lines. One T-DNA and two gene specific primers **(Table S1)** were used in three combinations for genotyping: The first reaction (1) contained two gene-specific primers, flanking the T-DNA insertion; the second (2) comprised the T-DNA left border primer and the upstream gene specific primer; the third reaction (3) contained the T-DNA left border primer and the downstream gene specific primer. **(C)** Phenotype of Arabidopsis wild-type (Col-0) and *ami1-2* plants grown for either 2 weeks on ½ MS plates or 4 weeks on soil. **(D)** IAM hydrolase activity in *ami1* T-DNA insertion mutants and AMI1 overexpressing AMI1ind lines. Crude extracts were incubated for 3 h at 30 °C with 5 mM IAM as substrate. The produced IAA was detected after separation by RP-HPLC. Given are means ± SE (n = 5). Similar results were obtained in two independent experiments. Analysis of **(E)** IAM and **(F)** free IAA levels. Two-week-old seedlings were used for the determination of IAA and IAM by LC-MS analysis. Three independent biological replicates were assessed. Quantification of free IAA levels in *ami1-1*, *ami1-2*, AMI1ind-1, and AMI1ind-2 relative to the wild type (Col-0). Given is the standard error of the mean (n = 9). (Student’s *t*-test; \**p* ≤ 0.05, \*\**p* ≤ 0.01, \*\*\**p* ≤ 0.0001).

The phenotypic alterations observed in the *ami1* alleles were only moderate. Areal plant parts showed a slight growth reduction, particularly when grown on soil for a longer period of time (**Fig. 4C**). A more detailed analysis of the root system morphology of *AMI1* mutants employing the WinRhizo™ image analysis software confirmed this observation and revealed a significantly reduced root branching and total root length and area, respectively, for both *ami1* alleles (**Fig. S2**). Although the phenotypic alterations were not pronounced, we asked the question whether the altered *AMI1* expression translates into detectable changes in the chemotype. To this end, we examined IAM hydrolase activity in the *ami1* alleles (**Fig. 4D**). In comparison with wt, the amidase activity in *ami1-1* and *ami1-2* was not null but rather reduced by approximately 45-50%, pointing towards the existence of an additional IAM hydrolase in Arabidopsis. Next, we analyzed changes in IAA and IAM contents by mass spectrometry. We found a correlation between the decreased IAM hydrolase activities in the *ami1* mutants and endogenous IAA and IAM levels. Both mutant alleles showed significantly reduced auxin contents, ranging 15 to 30% under the wt level alongside with significantly higher IAM contents (**Fig. 4E,F**).

Secondly, we generated mutants conditionally overexpressing *N*-terminally *c-myc* tagged *AMI1* cDNA. In these lines, the *c-myc:AMI1* construct was under the control of the β-estradiol inducible XVE transactivator (Zuo *et al.*, 2000). After screening numerous homozygous AMI1ind T_3_-lines, two independent lines, AMI1ind-1 and AMI1ind-2, with considerable transgene expression (**Fig. S3A**) and significantly increased amidase activity (**Fig. 4D**) were selected for all following experiments. In contrast to the *ami1* alleles, 2-week-old AMI1ind lines displayed significantly higher IAA levels and nearly 60% less IAM relative to wild-type plants (**Fig. 4E,F**). However, the root system architecture of the AMI1ind lines resembled the wt (**Fig. S2A-C**). To survey the influence of high-level *AMI1* expression on later stages of plant development, the AMI1ind mutants were grown on sterilized soil. Expression of the transgene was induced by irrigating the plants with water containing 50 μM β-estradiol (**Fig. S3A**). In comparison to non-induced plants, the induced AMI1ind plants exhibited an auxin-related phenotype, showing retarded growth (**Fig. S3B**) as well as curled leaf shapes (**Fig. S3C**), reminiscent of other plants showing auxin overproduction (Hentrich *et al.*, 2013; Zhang *et al.*, 2007; Zhao *et al.*, 2001). Furthermore, the AMI1ind lines showed considerably accelerated flowering when assessed on both a simple time and on a leaf number basis (**Fig. S3D**).

To further characterize the impact of *AMI1* overexpression, we monitored the root growth in response to the suggested substrate of AMI1, IAM. AMI1ind lines and wt plants were grown on plates containing rising amounts of IAM (**Fig. S4A**). With rising concentrations, the growth inhibiting effect of IAM became more pronounced in AMI1ind plants relative to the wt. At 200 μM IAM, the primary root growth was nearly fully suppressed in AMI1ind. Under these conditions, the overexpressors carried predominantly adventitious roots on their hypocotyls. In contrast, wt seedlings grown in parallel were at least able to produce short primary roots with an increased number of lateral and adventitious roots. The described adventitious root phenotype at high IAM levels is in line with the phenotype previously observed for tobacco plants constitutively overexpressing the *Agrobacterium tumefaciens iaaM* and *iaaH* genes (Sitbon *et al.*, 1992).

To quantify the observed differences, primary root elongation responses towards IAM of AMI1ind, *ami1*, and *35S::tms2* were determined and compared to wt Arabidopsis (**Fig. 5**). The *35S::tms2* mutant served as a control to assess the effect of *AMI1* overexpression. The line contains the 35S-driven *INDOLEACETAMIDE HYDROLASE (tms2*) gene from *Agrobacterium tumefaciens* (Karlin-Neumann *et al.*, 1991). Seedlings of the different genotypes were grown on ½ MS plates for 7 days, before they were transferred to plates containing IAM. While *ami1-1* and *ami1-2* seemed to be less sensitive towards IAM in the medium (**Fig. 5A**), AMI1ind-1 and AMI1ind-2 primary roots exhibited hypersensitivity towards IAM. Resembling the root elongation response of *35S::tms2*, both AMI1ind lines exhibited reduced root growth with rising IAM contents. At very high IAM concentrations (100 μM), however, root elongation was equally suppressed in all tested seedlings (**Fig. 5B**). The auxin response of the *AMI1* gain-of-function mutants was also examined in a similar manner employing plates containing different amounts of IAA (**Fig. S4B**). Only at low IAA contents (0.01 μM) AMI1ind and *35S::tms2* seedlings showed significantly less primary root elongation, which might be attributable to the higher intrinsic IAA level in those lines. With higher concentrations of IAA (0.1 to 10 μM) the growth response of the mutant roots was indistinguishable from those of the wt controls. Hence, it can be concluded that IAA perception is not affected by the conditional overexpression of *AMI1*. In summary, the metabolomic data underscore a contribution of AMI1 to auxin homeostasis. The only mild phenotype of the *ami1* mutants, however, implies that the role of the enzyme is likely not the general supply of auxin, but possibly the control of the cellular IAM pool.

**Figure 5.**
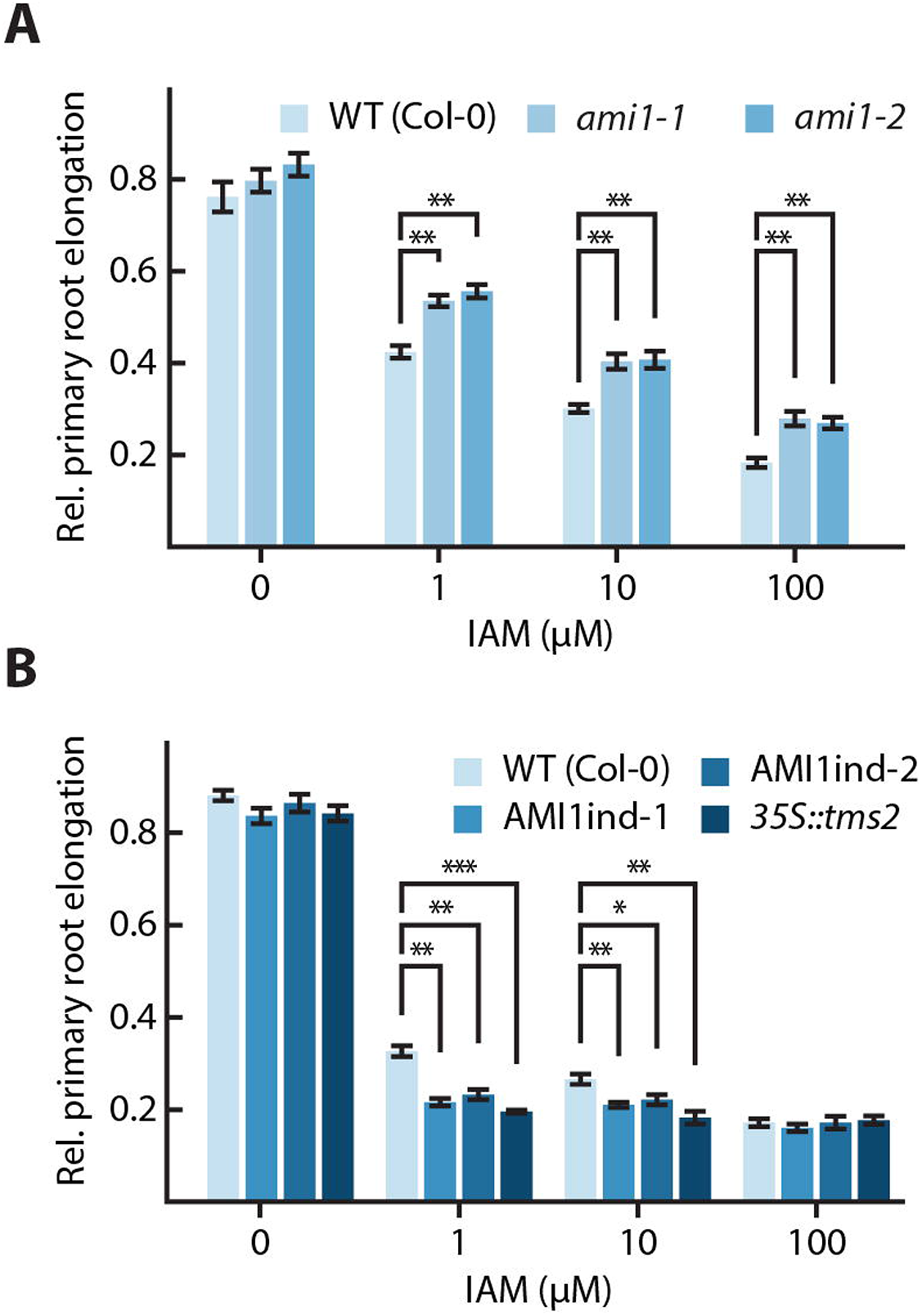
Root growth response of *ami1* and AMI1ind mutants towards IAM. Comparison of the root growth response of **(A)** *ami1* knockout and **(B)** IAM amidohydrolase overexpressing mutants with wild-type seedlings. Seeds were germinated and grown for 7 days on ½ strength MS plates, before they were transferred onto plates containing indicated amounts of either IAM or IAA. In order to examine the sensitivity toward IAM and IAA in the media without the bias of initially shorter primary roots, the impact of the two compounds was expressed in relative terms. Therefore, the length of the longest primary root of each genotype grown under control conditions was set to a value of 1 and all other roots of the corresponding genotype were expressed relative to this value. The primary root elongation of seedlings was quantified. The expression of the transgene in the AMI1ind lines was induced by adding 10 μM ß-estradiol to the plates. At least sixteen seedlings of each genotype were measured for each condition. The data represent means ± SE. Asterisks indicate significant differences between the corresponding WT control and the tested genotypes under the described conditions. (Student’s *t*-test; \**p* ≤ 0.05, \*\**p* ≤ 0.01).

### Altered abundance of salt-, osmotic-, and wounding-related transcripts in *ami1-2*

To obtain further information on the molecular alterations triggered by perturbations of *AMI1* gene expression, we conducted transcriptome profiling of sterilely grown 12-day-old wt, *ami1-2*, and AMI1ind-2 seedings. Herein, we were especially interested in transcriptional changes of genes previously associated with auxin-related processes. Hence, we first analyzed the differential expression of a sub-group of 128 genes, related with auxin biosynthesis, conjugation/deconjugation, degradation, transport, and signaling. The study revealed that only very few of the tested genes were differentially expressed in the mutants (**Table S2**). Except of the induction of *YUC8* and *ILL5/IAR3,* no other auxin homeostasis-related genes appeared to be differentially expressed in *ami1-2*. YUC8 is involved in auxin biosynthesis (Hentrich *et al.*, 2013), and ILL5/IAR3 are two highly similar IAA-amino acid hydrolases, specific for IAA-Leu and IAA-Phe (Davies *et al.*, 1999; LeClere *et al.*, 2002). The induction of *YUC8* and *ILL5/IAR3* could possibly conduce to compensate the loss of *AMI1* in auxin biosynthesis. On the other hand, *YUC8* and *ILL5/IAR3* have been associated with biotic stress responses (Hentrich *et al.*, 2013; Truman *et al.*, 2010), which could also suggest a link of AMI1 with plant stress-related processes. For AMI1ind-2, we found the auxin conjugation-related genes *UGT75D1* and *GH3.17* to be significantly induced. Together with the de-regulation of a number of auxin signaling and transport components, including *LAX2*, *PIN4*, *PIN5*, *IAA1*, *IAA12*, *IAA14*, *ARF6*, *ARF7*, and *ARF16*, this likely represents an answer to the overproduction of IAA in the conditional mutant.

Next, we undertook a detailed profiling approach to obtain a broader overview of the transcriptomic alterations in the *AMI1* mutants. Based on an adj. *p*-value of < 0.05 and an arbitrarily chosen fold-change value of ≥ 1.5, 391 and 969 DEGs were selected in *ami1-2* and AMI1ind-2, respectively (**Table S2**). From these genes, 287 and 347 were up- and 88 and 622 were down-regulated in *ami1-2* and AMI1ind-2 (**Fig. 6A**). From the DEGs, 289 and 775 genes, respectively, could be assigned to 178 and 273 significant GO terms for *ami1-2* and AMI1ind-2 (**Table S3**).

**Figure 6.**
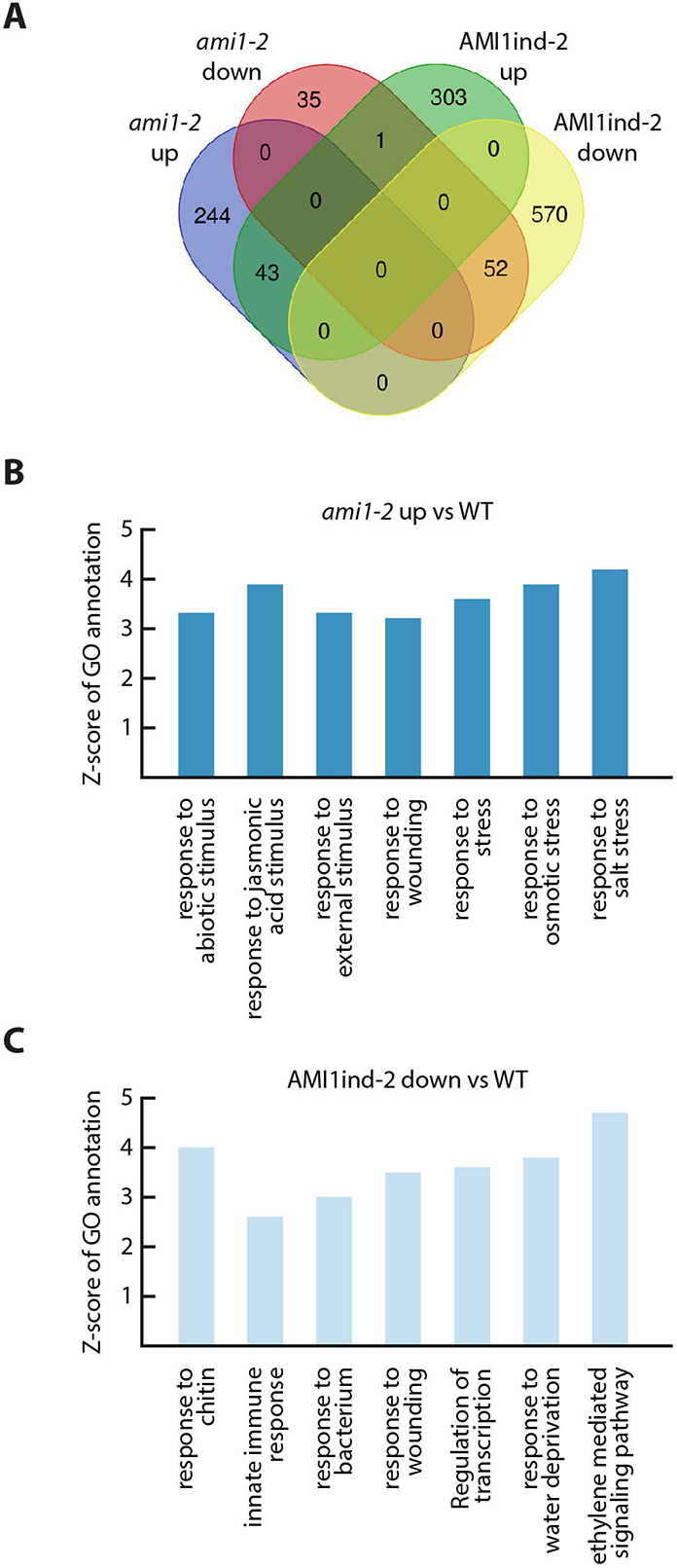
Transcriptome changes in ami1-2 and AMI1ind-2 are consistent with a contribution to plant stress responses. **(A)** Venn diagram visualizing the overlap between transcripts de-regulated in *ami1-2* and AMI1ind-2. Only genes showing a congruent directional change in transcript levels were scored as overlapping. **(B)** The top seven over-represented GO annotations for biological processes according to parametric analysis of gene set enrichment (PAGE) of the selected genes up-regulated in *ami1-2*. **(C)** The top seven under-represented GO annotations for biological processes according to PAGE of the selected genes down-regulated in AMI1ind-2. The Z scores show significant over/under-representation (Hochberg multiple testing adjustments in PAGE; *p* < 0.05). Fold changes of all entities were determined in 12-day-old *ami1-2* and AMI1ind-2 versus wild-type seedlings, respectively.

Parametric analysis of gene set enrichment (PAGE) suggested a prominent de-regulation of plant stress response-related genes in the *AMI1* mutants. Among the up-regulated genes in *ami1-2*, we observed the enrichment of genes related with salt- and osmotic stress, as well as oxylipin biosynthesis and signaling. The corresponding genes included, for instance, the AP2/ERF transcription factors *ERF53* and *RAP2.6*, the MYB transcription factors *MYB34*, *MYB47*, *MYB74*, *MYB102*, and *MYB108*, the JA signaling repressors *JAZ3* and *JAZ9*, *ALLENE OXIDE SYNTHASE* (*AOS*), *HYDROPEROXIDE LYASE* (*HPL1*), *JASMONIC ACID RESPONSE 1* (*JAR1*), the dehydration responsive genes *RD29*, *ERD7*, and *EDR9*, as well as the 9-*cis*-epoxycarotenoid dioxygenase gene *NCED3* (**Fig. 6B, and Fig. S5**). Among the down-regulated genes in AMI1ind-2, we found an over-representation of biotic- and drought stress-related genes. This second group of genes included, among others, the transcription factors *WRKY33*, *ORA47*, *ZAT10*, *DREB1B* and *DREB26*, as well as *PHENYL ALANINE AMMONIA LYASE 3* (*PAL3*), *MITOGEN ACTIVATED PROTEIN KINASE 3* (*MAPK3*), and the *RESPIRATORY BURST OXIDASE HOMOLOGUE D* (*RBOHD*) (**Fig. 6C, and Fig. S5**). Altogether, the transcriptome profiling implied a tight relationship of AMI1 with plant stress responses. Intriguingly, the genetic perturbations of *AMI1* expression had strong impact on gene expression regulatory processes. Pathway analysis using the MapMan tool (Thimm *et al.*, 2004) highlighted that AP2/ERF-, MYB-, and WRKY-class transcription factors were among the most affected molecular components associated with the regulation of transcription (**Table S3**).This likely suggests a role for IAM, or one of its derivatives, as a signaling molecule.

### The *ami1* mutants are hypersensitive to osmotic stress and accumulate ABA

Transcriptome profiling of *ami1-2* implicated a possible connection of IAM accumulation with abiotic stress responses. To gain additional insights into how AMI1 could be involved in plant stress responses, we examined the transcriptional response of *AMI1* towards different abiotic stress conditions (**Fig. 7**). While drought stress and salinity had either no or very little impact on *AMI1* expression, the osmotic stress treatment strongly repressed the expression of *AMI1* (**Fig. 7B**). To further characterize salinity and osmotic stress responses in *ami1*, we analyzed the growth behavior of mutant and wt seedlings under stress conditions. The survival rate of the seedlings has been assessed on the basis of first leaf emergence. When grown on 100 mM NaCl, *ami1* seedlings showed no considerable difference to wt. However, under osmotic stress conditions the survival rate of *ami1* seedlings was clearly reduced (**Fig. 7A**). The hypersensitivity of *ami1* seedlings towards osmotic stress conditions implies that the controlled repression of *AMI1* expression under osmotic stress conditions is an important mechanism in the stress adaptation process.

**Figure 7.**
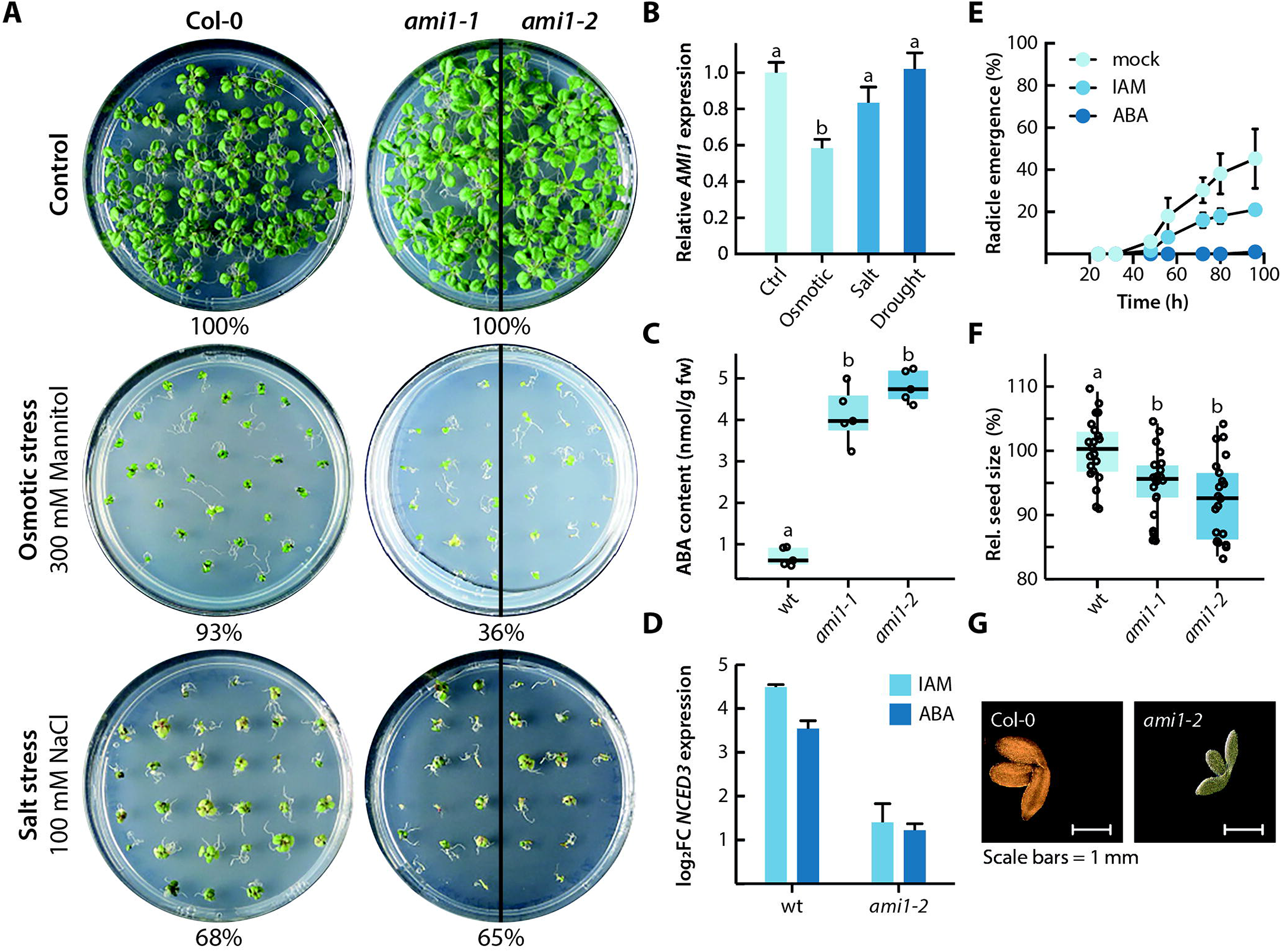
Relationship between *AMI1* expression levels and ABA. **(A)** Abiotic stress assay growing control plants (Col-0) and *ami1* mutants for 12 days under control, salt (100 mM NaCl), and osmotic stress (300 mM mannitol) conditions. Stress resistance was assayed on the basis of first true leaf establishment. Qualitative data are given in the figure. **(B)** Transcriptional changes in *AMI1* expression in response to different stress treatments in comparison to control conditions. Gene expression levels were normalized to the house-keeping gene *APT1* using the 2^−ΔΔC^T method. Given are the means with their SE (n = 9). **(C)** GC-MS/MS quantification of ABA contents in wild-type Arabidopsis (wt) and the two *ami1* mutant alleles. The box plots display the median, quartiles and extremes of the compared data sets (n = 5). **(D)** Induction of *NCED3* gene expression by 10 μM IAM and ABA in wild-type (wt) and *ami1-2* seedlings, respectively. Gene expression levels were normalized to the house-keeping gene *APT1* using the 2^−ΔΔC^T method. Given are the means with their SE (n = 9). **(E)** Effect of either 10 μM ABA or IAM on the germination of Arabidopsis seeds. The germination rate was analyzed on the basis of radicle emergence. Given are the means with their SE (n = 20). **(F)** Relative seed size of wild-type (wt) Arabidopsis and *ami1* mutant seeds. To analyze the seed size, seeds were photographed and the perimeter of at least 20 seeds determined using the Fiji software (Schindelin *et al.*, 2012). Different letters indicate significant differences between the means of the compared data sets analyzed by ANOVA and Tukey’s B post-hoc test (*p* < 0.05). **(G)** Size difference of representative wild-type (Col-0) and *ami1* mutant embryos. The photographs are scaled to the same size and scale bars (1 mm) are given.

Osmotic stress responses are mediated by ABA dependent and ABA independent pathways (Agarwal *et al.*, 2006; Yoshida *et al.*, 2014). The microarray data underpinned the activation of both pathways through the transcriptional regulation of key genes such as *NCED3* as well as several *MYB*- and *DREB* class transcription factor genes, respectively. Dioxygenases of the NCED family catalyze a rate-limiting step of ABA biosynthesis. Mass spectrometric assessment of ABA levels in wt and *ami1* mutants revealed significantly increased hormone levels in the mutants (**Fig. 7C**). To investigate whether *NECD3* expression is responsive to ABA or to IAM, the transcriptional response of the gene was analyzed by qPCR. Both substances induced *NCED3* expression (**Fig. 7D**). However, the observed effects were stronger in wt compared to mutant seedlings. Most interestingly, IAM showed a stronger effect in wt seedlings relative to ABA, which suggests that IAM acts either in parallel to ABA or upstream of the hormone in the transcriptional regulation of *NCED3*. Since *AMI1* gene expression was shown to be regulated by osmotic stress, we asked the question whether ABA could be involved in controlling *AMI1* expression in a feedback loop. On the basis of qPCR experiments and the quantification of promoter activity in *pAMI1::GUS* lines, we were not able to detect any regulatory effect of ABA on the expression of *AMI1* (**Fig. S6**).

### Impaired *AMI1* expression translates into decreased seed and embryo size

ABA is a major determinant of seed dormancy and germination (Vishal and Kumar, 2018). For this reason, we tested the effect of IAM on seed germination in comparison to ABA. Wild-type seeds were germinated either on ½ MS plates containing methanol (mock), or 10 μM IAM and ABA, respectively (**Fig. 7E**). As expected, ABA blocked seed germination nearly completely. Relative to the mock control, IAM also retarded the emergence of the radicle, most likely through the previously described induction of *NCED3* expression and the therewith coupled increase in ABA levels. A recent study reported the specific role of the KUP/HAK/KT-family K^+^ transporter KUP9 in controlling cellular auxin homeostasis through the symport of IAA from the eR into the cytoplasm (Zhang *et al.*, 2020). Work from our lab related IAA and IAM with potassium influx in Arabidopsis embryos through the transcriptional control of K^+^ transporter gene expression. The work also associated the two indolic compounds with the control of the seed size (Tenorio-Berrio *et al.*, 2018). In order to gain detailed information on the *ami1* mutant seed phenotype, we inspected seed- and embryo size of *ami1* loss-of-function mutants in comparison to wt (**Fig. 7F,G**). The seed size of both *ami1* mutant alleles was significantly reduced. Consistent with this finding, the embryos of the *ami1* mutants were also observed to be considerably smaller. Overall, the presented data suggest that IAM triggers ABA production through the stimulation of its biosynthesis. Furthermore, it can be concluded that the accumulation of IAM in the *ami1* mutants exerts a negative effect on seed maturation.

## DISCUSSION

### AMI1 contributes to auxin homeostasis *in vivo*

Auxins are well-characterized plant growth regulators ubiquitously distributed throughout the plant kingdom. Despite considerable scientific interest into this substance class, our knowledge of IAA biosynthesis and cellular homeostasis in plants is still incomplete. The formation of IAA is assumed to proceed via several independent pathways that either act redundantly or in parallel to each other (Pollmann *et al.*, 2006a; Woodward and Bartel, 2005; Zhao, 2010). However, only the major auxin biosynthesis pathway via indole-3-pyruvate is yet fully uncovered with respect to the enzymes and intermediates involved (Kasahara, 2016; Zhao, 2014). Among the discussed additional pathways, one is suggested to proceed via IAM that is further converted by IAM hydrolases to give rise to the active plant hormone, IAA. In recent years, evidence for the wide distribution of corresponding IAM hydrolases in the plant kingdom has been provided (Nemoto *et al.*, 2009; Pollmann *et al.*, 2003; Sánchez-Parra *et al.*, 2014), suggesting a conserved function of AMI1-like IAM hydrolases in plants. Here, we report the detailed characterization of *AMI1* loss- and gain-of-function mutants and show that AMI1 contributes to the conversion of IAM into IAA *in planta* (**Figs. 4-5** and **S2-4**). Remarkably, IAM hydrolase-activity was not null in *ami1-1* and *ami1-2* (**Fig. 4D**), which suggested the existence of other enzymes that might be involved in the conversion of IAM into IAA, but the analysis of the other members of the amidase signature (AS) family in Arabidopsis provided no indication for a second specific IAM hydrolase in this group (Neu *et al.*, 2007; Pollmann *et al.*, 2006b). A recent study, however, associated two additional formamidase-like proteins, IAMH1 and IAMH2, with this reaction (Gao *et al.*, 2020), which is likely the reason for the remaining enzymatic activity. It will be highly interesting to introgress the *ami1* mutation into the *iamh1 iamh2* CRISPR/Cas line in a future project, in order to fully understand the role of IAM in plants.

### The IAM-pathway is not essential for general IAA supply

The analysis of the root system architecture of *ami1-1* and *ami1-2* revealed significant differences in the specific root branch density, as well as the specific root area and length (**Fig. S2**). *AMI1* expression in primary roots declined shortly after germination (**Fig. 2**), but apparently a tight regulation of the spatio-temporal distribution of AMI1 is important to facilitate proper root growth. On the contrary, the conditional overexpression of AMI1 had no significant effect on root growth (**Fig. S2**). Taking the hypersensitive response of AMI1ind lines towards exogenously applied IAM into account (**Fig. 5**), this implies that the IAM pool size in roots is low or not accessible for recombinant AMI1. Our observation is consistent with the lack of any considerable root phenotype when the *iaaM* and *iaaH* gene from *Agrobacterium tumefaciens* are expressed under the control of their natural promotors in Arabidopsis (van der Graaff *et al.*, 2003). Apparently, the IAM pool in areal plant parts must be bigger or more accessible to recombinant AMI1, as the leaves of AMI1ind plants showed clear phenotypic alterations (**Fig. S3**) that can be associated with auxin overproduction, resembling the phenotypes of mutants such as *yuc1-D* (Zhao *et al.*, 2001) or the *sur2* mutant (Delarue *et al.*, 1998), although the effects in the AMI1ind lines are somewhat weaker.

However, on a whole plant-scale, the phenotypic alterations caused by mutations in *AMI1* are only moderate (**Fig. 4**, **Fig. S2**), and the transcriptomics analysis of *ami1-2* provided neither evidence for a consistent induction of genes involved in other auxin biosynthesis pathways, nor are *IAMH1* or *IAMH2* induced to compensate the lack of AMI1 in the mutant (**Tab. S2**). Hence, it must be concluded that the IAM pathway is not strictly necessary for the general supply of IAA in Arabidopsis, although some particular developmental processes, such as lateral root growth or seed maturation (**Fig. 7**), may depend on AMI1 action to achieve full effectiveness.

### Transcriptomics analyses revealed a link between AMI1 and plant stress responses

Our expression analyses demonstrated that *AMI1* transcription is differentially regulated by IAM and IAA, even though IAA appeared to be less effective controlling *AMI1* expression (**Fig. 3**). This further suggested a tight transcriptional connection between AMI1 and both its substrate and reaction product. Most interesting, however, was the observation that the genetic reduction of AMI1 levels caused no major alterations of auxin homeostasis-related gene expression. This contrasts with the results obtained for the conditional *AMI1* overexpression line. Here, the induction of a few auxin conjugation-related genes suggested the activation of compensatory effects, likely triggered to counteract auxin overproduction in the line (**Table S2**).

The comprehensive profiling of transcriptional alterations in *AMI1* mutants revealed a strong impact on genes related with biotic and abiotic stress responses, including key enzymes for the biosynthesis of JA and ABA (**Fig. 6, Tabs. S2-S3**). Along with numerous marker genes for ABA signaling, including e.g. *RD26*, *COR47*, *ERD4*, *ERD9* and *ERD10*, the gene for a key-enzyme of ABA synthesis, *NCED3*, appeared strongly induced in *ami1-2*. The overexpression of *NCED3* has recently been associated with an improvement of drought tolerance in soybean (Correa Molinari *et al.*, 2020) Both the induction of *NCED3* through IAM application and the accumulation ABA in the *ami1* alleles could be confirmed (**Fig. 7C,D**), which provided strong arguments for an intimate relationship between AMI1 and IAM contents, respectively, and ABA. The considerable induction of AP2/ERF transcription factors in *ami1-2* (**Table S3**) has to be particularly emphasized, as it underpins our observations of a close connection between AMI1 action and abiotic stress responses. The particular role of AP2/ERF transcription factor networks in orchestrating hormone and abiotic stress responses is well documented (Xie *et al*., 2019). Quantitative gene expression analysis revealed a significantly suppression of *AMI1* transcription by osmotic stress (**Fig. 7B**), and *ami1* mutants grown under osmotic stress conditions show a considerably increased sensitivity to osmotic stress conditions (**Fig. 7A**). In consequence, we conclude that the suppression of *AMI1* expression by osmotic stress and the therewith linked accumulation of IAM are important variables that contribute to the finetuning of ABA-governed stress response by triggering NCED3-mediated ABA biosynthesis in Arabidopsis.

ABA and gibberellins are essential determinants of seed development and dormancy (Carrera-Castaño *et al.*, 2020). A recent publication also highlighted a direct role of auxin in seed dormancy (Matilla, 2020). In addition, crosstalk between IAA and ABA in seed development and germination has been reported to occur on both the transcriptional and metabolic level (Chauffour *et al.*, 2019; He *et al.*, 2020). With respect to our data on seed development and germination (**Fig. 7E-G**), it can be concluded that the crosstalk between AMI1/IAM contents and ABA-related processes is also involved in facilitating proper seedling development and germination. We were able to demonstrate that IAM application reduces the germination rate in Arabidopsis, likely through the induction of ABA biosynthesis. Moreover, mutations in *AMI1* translated into aberrant embryo and seed size.

The Arabidopsis AS superfamily member FATTY ACID AMIDE HYDROLASE (FAAH) is associated with numerous developmental processes, including plant growth and early flowering, and interacts with ABA signaling (Teaster *et al.*, 2012; Teaster *et al.*, 2007; Wang *et al.*, 2006). FAAH catalyzes the hydrolysis of *N*-acylethanolamines (NAEs), lipid signaling molecules that orchestrate a wide array of physiological processes in multicellular eucaryotes (Chapman, 2004), which terminates their action (Aziz and Chapman, 2020). Although there are marked differences between FAAH and AMI1 with respect to their substrate preferences (Pollmann *et al.*, 2006b), the growth inhibiting properties of their preferred substrates led us to the hypothesis that the main role of AMI1 might be similar to that of FAAH, i.e. terminating the action of IAM through its conversion to free IAA and NH ^+^. Particularly in plant stress responses and seed development, this mechanism involving the IAM induced biosynthesis of ABA (**Fig. 8**) seems to be important to drive proper adaptational responses. In consequence, the AMI1-dependent metabolic flux through the IAM shunt is seemingly an important regulatory variable that connects auxin-mediated plant growth processes with plant stress responses. It will be highly interesting to further investigate the regulatory role of IAM in plants and how the corresponding signal is perceived and translated into downstream responses. At the same time, it appears tempting to us to further investigate the transcriptional networks that are involved in the integration of the IAM signal.

**Figure 8.**
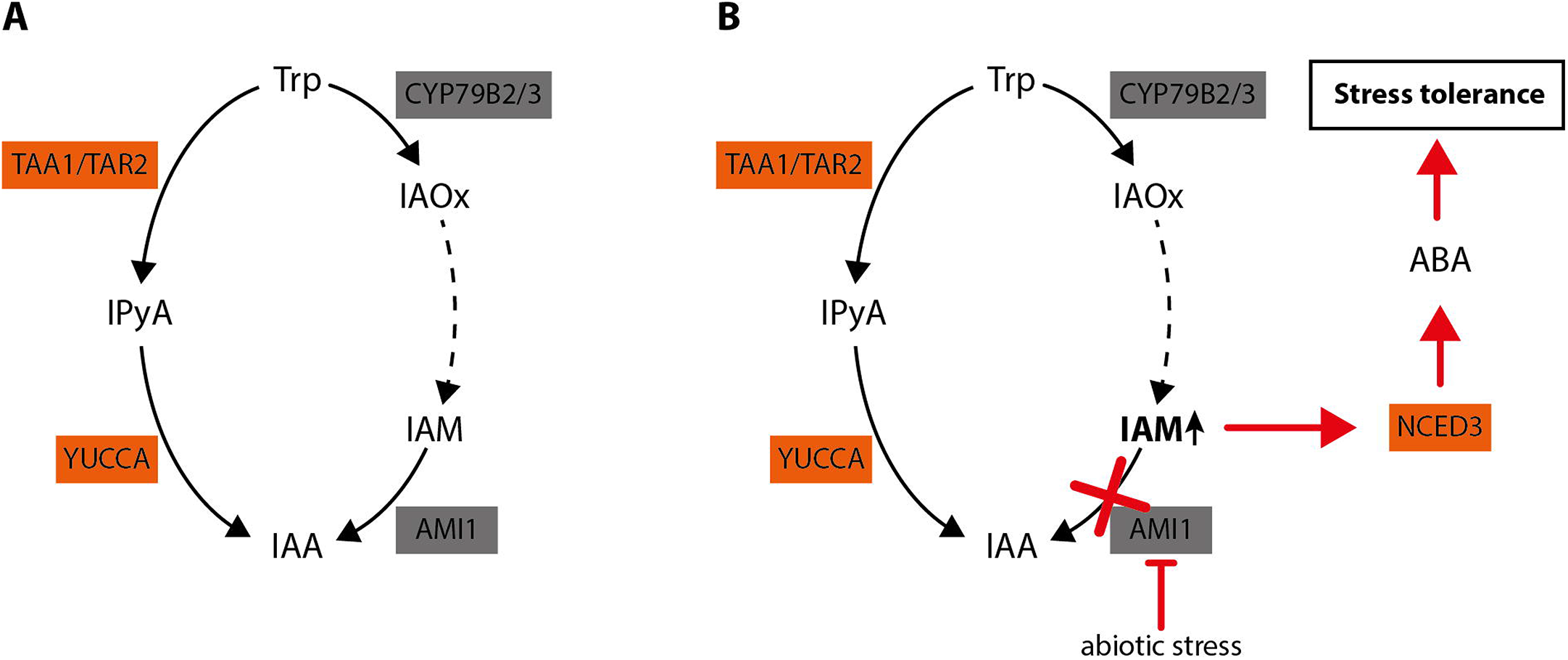
Integrated model of the molecular role of AMI1. **(A)** Our data suggest that the metabolic flux of the parallel pathways normally leads without distortions to IAA. In this scenario, IAM is not considerably accumulating as it is quickly converted into IAA. **(B)** Under stress conditions, however, the expression of *AMI1* is repressed. In consequence, this results in the accumulation of IAM. The increasing IAM levels gradually induce the expression of *NCED3* and trigger the subsequent biosynthesis of ABA as well as the corresponding downstream processes that confer stress tolerance in Arabidopsis.

## ABBREVIATIONS

RP-HPLC: reverse-phase high performance liquid chromatography
LC-MS: liquid chromatography-mass spectrometry
GC-MS: gas chromatography-mass spectrometry
IAA: indole-3-acetic acid
IAM: indole-3-acetamide
IAOx: indole-3-acetaldoxime
IPyA: indole-3-pyruvic acid
AMI: amidase

## ACKNOWLEDGEMENTS

The authors thank the NASC for providing seeds of T-DNA insertion and mutant lines, and Prof. Nam-Hai Chua (The Rockefeller University, New York) for providing the pMDC7 vector. This research was supported by grants from the German Research Foundation (DFG, SFB480/A10) and the Spanish Ministry of Economy, Industry and Competitiveness (MINECO, BFU2017-82826-R to SP and a grant from the Swedish Research Council (VR) to HA.

## Supplementary data

**Protocol S1.** Supplementary experimental procedures.

**Table S1.** Primers used for genotyping, cloning and expression analysis.

**Table S2.** Microarray analysis.

**Table S3.** GO and MapMan analysis of selected up- and down-regulated genes in the *AMI1* mutants.

**Figure S1.** Expression analysis of *AMI1* in wt and genotyping of *ami1* alleles.

**Figure S2.** Root phenotype of *AMI1* mutant plants.

**Figure S3.** Phenotype of conditionally *AMI1* overexpressing AMI1ind lines.

**Figure S4.** Root growth responses of conditional *AMI1* overexpression lines towards IAM and IAA in the media.

**Figure S5.** Validation of Microarray Data by qRT-PCR, related to Figure 6.

**Figure S6.** Effect of ABA on *AMI1* gene expression.

